# Digital holography-based 3D particle localisation for single molecule tweezer techniques

**DOI:** 10.1101/2022.05.03.490423

**Authors:** James L. Flewellen, Sophie Minoughan, Isabel Llorente Garcia, Pavel Tolar

## Abstract

We present a three-dimensional imaging technique for fast tracking of microscopic objects in a fluid environment. Our technique couples digital holographic microscopy with three-dimensional localisation via parabolic masking. Compared with existing approaches, our method reconstructs 3D volumes from single-plane images, which greatly simplifies image acquisition, reduces the demand on microscope hardware, and facilitates tracking higher densities of microscopic particles while maintaining similar levels of precision. We demonstrate utility of this method in magnetic tweezer experiments, opening their use to multiplexed single-molecule force spectroscopy assays. We propose that our technique will also be useful in other applications that involve the tracking of microscopic objects in three dimensions.

**SIGNIFICANCE:** Tracking objects in 3D is a common task in biology, but typically requires the acquisition of image stacks, which is limited by speed, the depth of field of microscope objectives and by the presence of other objects that obscure the illumination. Here we develop HoloMiP (<u>Holog</u>raphic <u>Mi</u>croscopy with <u>P</u>arabolic masking), which uses digital holography to reconstruct the three-dimensional images from a single plane allowing tracking of light-scattering objects in 3D. HoloMiP outperforms existing methods in precision, speed, simplicity and tolerance to crowding. We show that it is particularly suitable for fast, multiplexed magnetic tweezer experiments, opening new avenues to high-throughput force spectroscopy.

## INTRODUCTION

Tracking microscopic objects in a fluid environment is common in biology. It is used in magnetic and optical tweezers (1), in microfluidic devices (2), for quantifying fluid flows around cells (3) and for direct analysis of the motion of particles, microorganisms, or cells (4). While two-dimensional (2D) tracking in the image (xy) plane is well-established, many applications call for information along the third dimension, parallel with the optical axis (z). Three-dimensional (3D) tracking is challenging because capturing the motion of objects through the imaged volume typically requires acquisition of z-resolved image stacks, which severely limits time resolution. For single light-scattering objects, continuous 3D acquisition is facilitated by the use of look-up table techniques, which infer the z position by matching interference rings around the object to reference image stacks (5). However, such techniques require access to each tracked particle prior to or post-tracking to acquire reference z-stacks and are limited by the depth of field of microscope objectives and by the presence of other objects in the sample, which obscure the illumination source.

Magnetic tweezers (MT) is a typical imaging application that relies on 3D tracking (6). Magnetic tweezers uses an external magnetic field to apply a force or torque to microscopic superparamagnetic beads, which are conjugated to molecules of interest. The position of these microbeads is tracked through time in order to monitor the effects of the external force. This technique has traditionally been used to study force- or torsion-dependent molecular processes, for instance: the extension and torsion of DNA (7, 8), the action of helicases and other DNA-binding proteins (9), protein unfolding (10), and force-dependent proteolysis (11). However, the technique also shows promise in probing the kinetics of single-molecule ligand binding at low applied forces and with high throughput (6,12–14). The full potential of MT in this application, however, has not yet been realised due to difficulties arising from the uncertainty about the number of interacting molecules, the non-specific interactions of the microbeads with the imaging surface (15), and also the challenge of tracking the position of the microbeads in three dimensions with sufficient time resolution.

Standard magnetic tweezer protocols localise magnetic microbeads along z by taking advantage of diffraction rings emanating from defocused beads when a coherent illumination source is used. Because the pattern of the diffraction rings depends on the microbead distance from the objective focal plane, the z position of the microbead can be determined by comparing its diffraction pattern to that of a previously determined look-up table (5). The look-up table is a z-stack of microbead images recorded by capturing the diffraction rings at known z positions, typically using a nano-positioning stage, whilst an external magnetic field is applied to lift the microbeads from the surface of the imaging chamber. However, the need to acquire a look-up table for every microbead that needs to be localised severely limits this technique in cases when microbeads move or dissociate during the experiment due to the applied external magnetic field. The cross-correlation algorithm (5, 16), which is used to compare the ring pattern to the look-up table, also depends on symmetry and fails when diffraction patterns overlap. This restricts the density of objects that can be tracked in a single field of view. Finally, not all microscopes have a stage with sufficient nano-positioning precision for this technique.

Our technique uses inline digital holographic microscopy (17) to generate a three-dimensional reconstruction of the light field from a single plane image. Objects are localised in this field as peaks of intensity using a 3D parabolic masking technique. We term our technique HoloMiP (<u>Holo</u>graphic <u>Mi</u>croscopy with <u>P</u>arabolic masking). HoloMiP does not require generation of look-up tables or any *a priori* knowledge of the objects being imaged and it can localise them even if their diffraction patterns overlap. In addition, its application is not reliant on the symmetry of the imaged objects, and could be readily applied to non-spherical microscopic objects that scatter light, such as microflakes (18) or rod-shaped bacteria (19). We demonstrate the utility of HoloMiP in fast, multiplexed single molecule force spectroscopy assays using MT to apply force at receptor-ligand bonds attached to a surface by DNA tethers.

## RESULTS

### HoloMiP algorithm

To develop HoloMiP for three-dimensional particle tracking, we used an inverted microscope coupled with magnetic tweezers similarly as described (20). The sample containing superparamagnetic microbeads tethered to the coverslip is illuminated by a semi-coherent light-emitting diode and imaged with a high-magnification objective focused above the sample. In order to control the vertical position of the tethered beads, a pair of permanent magnets move vertically above the sample using a fast voice-coil actuator.

The semi-coherent illumination produces diffraction patterns of the microbeads on the xy image plane (Fig. 1a), which are treated as holograms to recover the 3D positions of the beads (21). We use the Rayleigh-Sommerfeld formalism (22) to reconstruct the 3D electromagnetic field of the illuminated field of view, in a step termed ‘numerical back-propagation’ (23) (Fig. 1b, c; Supplementary Fig. 1a-c and Supplementary videos 1–6, see Methods for the full description of the procedure). Volumes of high electromagnetic intensity are used as proxies for the locations of the microbeads. Subsequent computational steps interrogate the reconstructed 3D electromagnetic field to localise precisely the positions of peaks in the high-intensity volumes. Approximate transverse (x, y) coordinates of each microscopic object are found by first projecting the maximum-intensity pixel in each transverse plane to a single plane, then deploying a 2D peak-finding algorithm (Fig. 1d). Narrow cuboids along the optical axis centred on these positions are examined (Fig. 1e) and a Sobel-like gradient filter (24) is applied to the field in these cuboids (Fig. 1f; Supplementary Fig. 1d-f, Supplementary Fig. 2 and Supplementary videos 4-7) to identify an approximate z position of each object. A small cuboid around each initial x, y, z object position (Fig. 1g) is then used for precise 3D localisation using a 3D version of parabolic masking (25) (Fig. 1h-j). Detailed information on the localization approach can be found in Methods.

**Figure 1.**
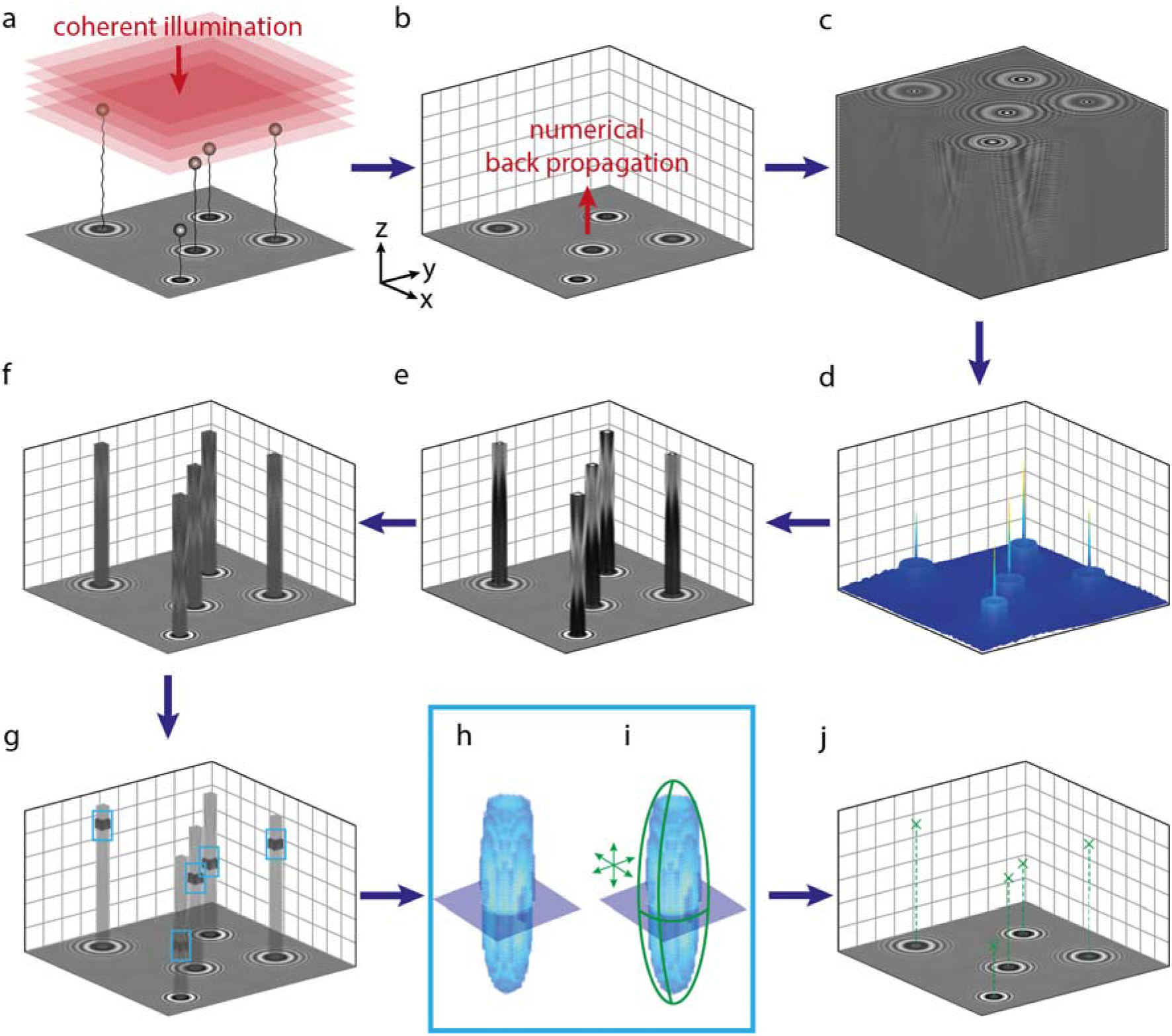
HoloMiP algorithm for localising microscopic objects in 3D. (a) Sample is illuminated by a coherent light source, resulting in a recorded digital hologram. (b) Rayleigh-Sommerfeld back propagation reconstructs the electromagnetic field at each user-defined plane, resulting in (c) a 3D electromagnetic field over the sample volume. (d) an initial guess for the x, y position of each object is found by taking a maximum projection through the intensity of the electromagnetic field, followed by a 2D peak-finding algorithm. (e) Columnar cuboids along z and centred on each initial x, y position are extracted. (f) a 3D Sobel-like gradient filter is applied to these cuboids. An initial guess for the z position is found as the maximum intensity along z. (g) Cuboids around the initial x, y, z guesses are extracted and used for precise localisation by parabolic masking; (h) shows an isosurface representation of one of the objects. (i) A 3D parabolic surface is moved around the cuboid until it matches the data. This results in a precise 3D position for the objects (j) which can be used for further analysis.

The reconstructed z position is corrected to account for the mismatch in refractive indices between the immersion oil and the sample medium, which leads to an underestimation of the reconstructed z displacement of an object (26). The application of this optical correction results in the retrieval of accurate absolute particle positions. While a correction factor of the ratio between refractive indices based on the small-angle approximation is commonly used (27), we take inspiration from another approach (16) and develop an empirical correction factor. We used a nano-positioning piezo stage to scan immobile microbeads along the optical axis and reconstruct their positions using HoloMiP. We found that it is possible to apply a linear correction to reconstructed z positions ranging between 7 and 12 μm from the focal plane for a 40× objective to recover the absolute positions. Outside of this range, a non-linear correction factor can be applied (see Supplementary Material and Supplementary Fig. 3).

To characterise the potential computational burden of HoloMiP, we measured processing times on a dedicated workstation with an Intel i9 CPU with 14 cores and with 128 GB total RAM. With this setup, a typical hologram frame 1024 px × 1024 px and reconstructed to 100 z-slices takes ^~^1.1 seconds to output the 3D positions of all objects. The computational demands are manageable in typical experiments. Furthermore, this time is virtually independent of the number of objects to be localised in the field of view. By contrast, the computational time of the LUT method is heavily dependent on the number of objects in the field of view. Thus, while HoloMiP could be more demanding of computational power than the LUT technique for low densities of objects, at higher densities it shows notable benefits. See Methods for more details.

### Comparison between HoloMiP and cross-correlation look-up table techniques for 3D localisation of microbeads

We compared the performance of HoloMiP with the cross-correlation look-up table method used commonly in MT experiments (16). To compare x, y localisation, a synthetic diffraction profile of a defocused microbead was generated *in silico* and moved in-plane in discrete steps over a background image (Fig. 2a-c). The size of the steps was adjusted from 0.3 nm to 19.5 nm and the position of the microbead was determined using both HoloMiP and a cross-correlation look-up table technique. The resolving performance of the two techniques was compared by calculating the signal-to-noise ratio (SNR) as the step size divided by the mean standard deviation of the position trace throughout the recording (Fig. 2d). HoloMiP had consistently higher SNR than the look-up table technique due to lower mean standard deviation (0.68 nm for HoloMiP versus 1.63 nm for the LUT). Consistently, the noise value approached the signal around a step size of 0.7 nm for HoloMiP, whereas for the look-up table technique, the value was around 1.6 nm. This analysis indicates that under matching conditions HoloMiP is capable of detecting smaller lateral displacements with higher SNR compared to the look-up table technique.

**Figure 2.**
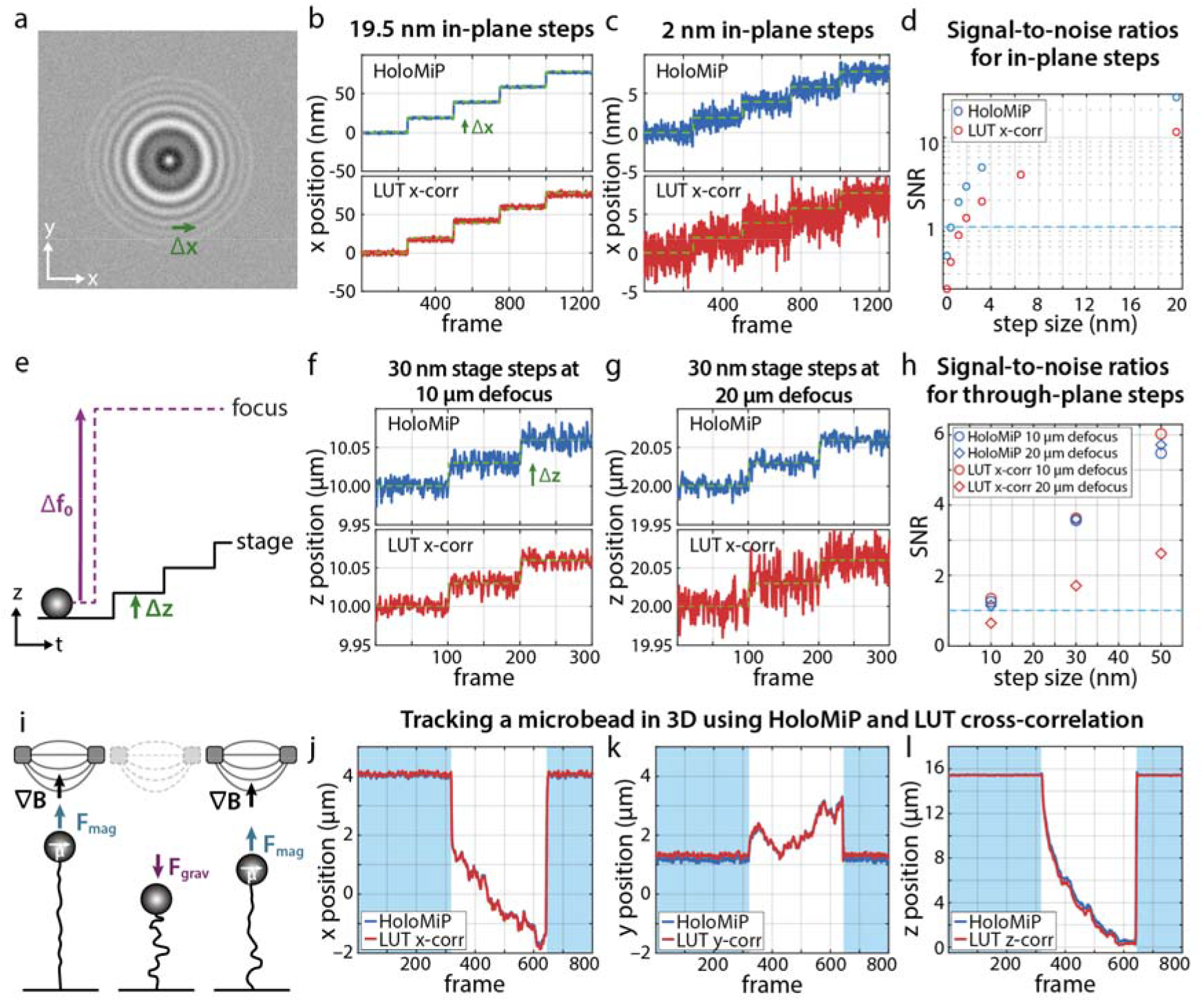
Comparison of HoloMiP and look-up table cross-correlation (LUT x-corr) for localising superparamagnetic microbeads. (a) An *in silico* synthetic diffraction pattern, positioned 10 μm from the focus, is moved in-plane by discrete steps (Δx). The x, y localisation through time of the diffraction pattern moving in steps of 19.5 nm (b) and 2 nm (c) are shown for each technique. The mean standard deviation was 0.68 nm for HoloMiP and 1.63 nm for the LUT. (d) The signal-to-noise ratios (SNR) are shown for each technique. The dashed line shows a SNR value of 1. (e) The through-plane resolution of the two techniques is tested by using a nano-positioning stage to move an immobilised microbead in steps (Δz) along the optical axis (z) after an initial defocusing of the microscope objective (Δf_0_). Resolution of 30 nm steps with a 10 μm (f) and 20 μm defocus (g). (h) The corresponding SNR shows the two techniques are similar at a 10 μm defocus; however, HoloMiP outperforms the look-up table technique at the 20 μm defocus. (i) The magnetic field gradient (**∇B**) from two permanent magnets induces a magnetic dipole (**μ**) in, and exerts an upwards force (**F**_mag_) on, a 2.8 μm magnetic microbead tethered to a 16.3 μm length of DNA. When the magnets are moved out of position, the microbead descends to the surface under gravity (**F**_grav_). (j, k, l) The position of the microbead in response to the magnetic field is tracked through time in 3D using both HoloMiP and the look-up table technique; blue shading indicates when a magnetic force of 15 pN is applied.

To test the localisation precision along the optical axis (z) along with x and y, we immobilised microbeads on a glass surface and recorded images at 100× magnification with defocus of 10 μm or 20 μm from the sample plane. A nano-positioning stage then moved the bead sample along the optical axis in discrete steps of 10 nm, 30 nm and 50 nm (Fig. 2e-g). Although the localisation precision in x and y was lower than in the *in silico* experiments, the two techniques performed similarly at 10 μm defocus; however, at 20 μm defocus, HoloMiP outperformed the look-up table method in z-precision: at 10 μm defocus, the mean standard deviation along x, y and z was 5.2 nm, 7.7 nm and 8.0 nm, respectively, for the look-up table technique, and 5.6 nm, 9.9 nm and 8.6 nm for HoloMiP. At 20 μm defocus, the corresponding values for HoloMiP were comparable to 10 μm defocus along all three directions (4.4 nm for x, 8.1 nm for y, 8.7 nm for z), however, for the look-up table technique, the values were only comparable along × (4.2 nm) and y (6.0 nm), but a factor of two greater along z (17.6 nm). This was reflected in differences in SNR calculated as above (Fig. 2h).

Thus, we conclude that HoloMiP matches the look-up table technique in z position precision and extends the focus range through which objects can be tracked with high precision.

To compare directly the two techniques in a MT experiment, we analysed the 3D movement of a 2.8 μm-diameter Dynabead, attached to the surface via a DNA tether, in response to an applied magnetic field (Fig. 2i-l). The bead is tethered to a 16.3 μm length of λ-DNA via multiple biotin-streptavidin attachments at the glass surface and digoxigenin–anti-digoxigenin at the bead. At the start of the trace, the DNA tether is fully extended through a force of 15 pN induced by the magnets acting on the bead. The magnets are then removed, and the position of the microbead is tracked as it sediments to the floor of the chamber. The magnets are then reintroduced, and the bead lifts away from the surface, re-extending the tether. The outputs of HoloMiP and the look-up table technique were in close agreement, although not identical. We noted a lateral offset between the two techniques, which increases linearly with microbead distance from the focus (Supplementary Fig. 4) and is independent of bead position in microscope field of view. We hypothesise that HoloMiP is sensitive to the alignment of the optical axis to the camera sensor in a way that the LUT technique is not. The LUT technique involves averaging a 2D diffraction pattern to produce a 1D radial profile to be compared to a look-up table. This method imposes a radial symmetry on the diffraction pattern, removing any eccentricity in the diffraction ring patterns. Elliptical ring patterns would occur when the optical axis is not perfectly normal to the imaging plane. This averaging does not occur with the holographic reconstruction in HoloMiP, resulting in a sensitivity to an angle between the normal of the imaging plane and the incident illumination. It is important to note firstly, that this effect is small and secondly, that it does not affect the determination of *relative* displacement of objects through time, which is used to determine relevant biophysical parameters. If desired, the two approaches can be made to agree using a simple linear correction factor (see Supplementary Material).

### HoloMiP is able to localise high densities of particles

A weakness of the cross-correlation look-up table technique is a tendency to fail when the diffraction patterns from adjacent objects overlap. To assess how HoloMiP performs in localising particles with overlapping diffraction patterns, we compared the two techniques in experiments with *in silico* synthetic diffraction patterns of microbeads. First, we performed a ‘crowding’ experiment, whereby a central microbead was surrounded by up to eight ‘crowding’ beads in the same xy plane and at distances ranging from 8 to 24 μm, corresponding to ^~^3-8 bead diameters (Fig. 3a-c). We recovered the z position of the central bead in all conditions as the distance from the focal plane varied from 0 to 30 μm using HoloMiP or the look-up table technique and compared them to a baseline position determined from images containing the central bead only (Fig. 3d, e). As expected, the look-up-table technique failed when bead separation approached 10 μm (^~^3.6 bead diameters), especially at greater focal distances, where the diffraction rings are larger (see white crosses in Fig. 3d, e). By contrast, HoloMiP was able to recover the localisation of the central bead in all conditions, up to 30 μm defocus. As an indication of localisation precision, the magnitude of the greatest difference between the baseline bead location and HoloMiP’s localisation was ^~^0.6 μm along z (approximately one-fifth of the microbead diameter) and ^~^0.04 μm within the image plane. Overall, these results show a significant advantage of HoloMiP with respect to look-up-table techniques for the localisation of closely spaced diffracting microbeads.

**Figure 3.**
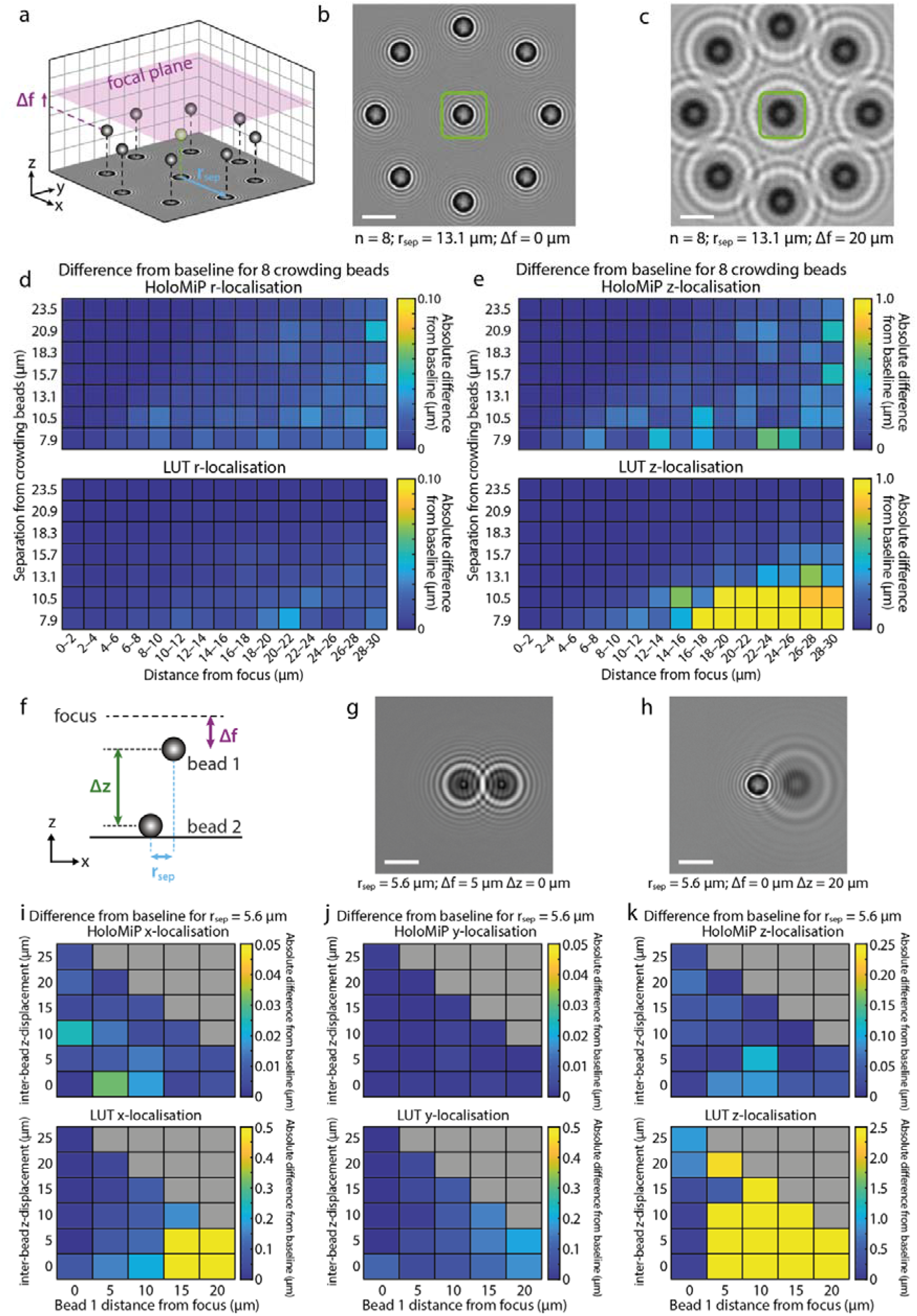
HoloMiP is successful at localising microbeads in 3D when their diffraction rings overlap. (a) Schematic for an *in silico* synthetic diffraction pattern bead ‘crowding’ experiment. Up to eight microbeads (n) surround a central bead (in green) at a range of separation distances (r_sep_). The plane containing the microbeads is displaced from the focal plane by Δf, which ranges from 0 to 30 μm. (b) and (c) show two example fields of view for n = 8. The 3D position of the central bead is recovered using both HoloMiP and the look-up table technique and compared to the positions found for a baseline image with no crowding. (d) and (e) show variation from this baseline for the mean of 2 μm intervals of Δf for both in-plane 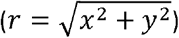 and z localisations. Note that the look-up table technique is less successful at z-localisation at greater focus distances. (f) Schematic for a synthetic diffraction pattern bead experiment where a single crowding bead (bead 2) is positioned closely to the central bead (bead 1) along x, but the z-displacement (Δz) between the beads is varied (g) and (h). (i), (j) and (k) show the effect of the crowding bead on the precision of localisation of the central bead, when compared to a baseline value with no crowding bead. Note that the scale on the LUT plots is an order of magnitude different to that on the HoloMiP plots. Grey squares indicate experimental conditions not tested. Scale bars are 5 μm.

Second, we investigated the effect of crowding beads co-localised closely in x and y, but displaced along z. In *in silico* synthetic diffraction pattern bead experiments with two beads, we varied the position of the central bead (bead 1) from the focus, and the z distance of the crowding bead (bead 2) from the first bead (Fig. 3f-h). We also assessed effects of such crowding when the crowding bead was displaced in x by one to two bead diameters, which is a realistic scenario in MT assays. Again, the look-up table failed to recover the localisation of the bead of interest (see white crosses in Fig. 3i-k), except in some cases where the central bead was sharply in focus, while HoloMiP was able to recover the localisation of the central bead in all conditions tested (Fig. 3g, h). As an indication of localisation precision, the greatest difference between the baseline bead position and HoloMiP’s localised position along x was ^~^0.03 μm when the microbeads were separated by two bead diameters, and ^~^0.05 μm with a single bead diameter displacement. The corresponding values along z were ^~^0.11 μm and ^~^0.23 μm. Effects along y were negligible.

### Measurement of force-mediated dissociation of antibody-antigen bonds using HoloMiP

To demonstrate the advantages of HoloMiP for a new MT application, we designed a system to measure dynamic single-molecule bond dissociation. Using a look-up table in such applications limits the number of microbeads that can be analysed in the field of view and prevents measurement of fast dissociation because of the need to acquire a z stack prior to each measurement. Our experimental system consisted of antibody-conjugated superparamagnetic microbeads, bound to antigen covalently attached to a DNA tether, in turn bound to the surface of the imaging coverslip via multivalent streptavidin-biotin interactions (Fig. 4a). The application of magnetic forces extends the DNA tether before bond dissociation, thus separating the antibody-antigen interaction from the chamber surface. This has two advantages. First, it eliminates binding contributions from non-specific interaction between the antibodies or microbeads with the surface. Second, by measuring the extension length of the DNA tethers before dissociation, beads tethered through multivalent interactions can be eliminated from the kinetic analysis of dissociation as they show shorter tether extension (Supplementary Fig. 5a).

**Figure 4.**
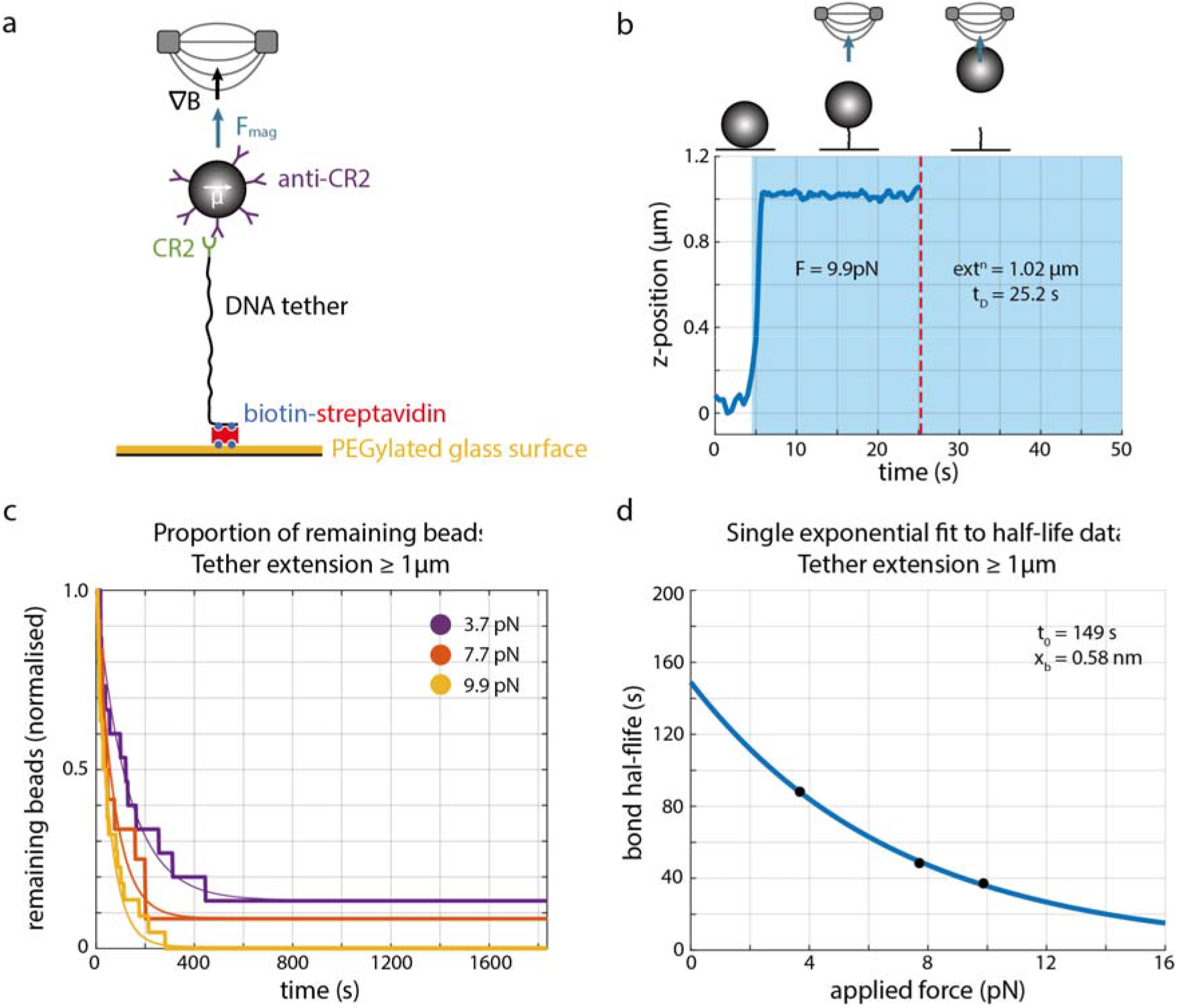
Application of HoloMiP to magnetic tweezer assays studying force-mediated single-molecule dissociation. (a) Experimental setup to probe dissociation between CR2 and anti-CR2 antibodies. (b) Example z-trace of a tethered microbead subject to a 9.9 pN force. Blue shading indicates when the magnetic field is applied; dissociation time is taken when the microbead disappears from the field of view (red line). Images are acquired at 2 fps before the magnetic field is applied; 5 fps for the first 30 seconds after the magnetic field application and 2 fps for the duration of the recording. (c) Proportion of microbeads, which extended to at least 1.0 μm from the chamber surface, remaining attached in response to three different applied forces. Single exponentials have been fitted to determine k_off_ values (F = 3.7 pN, N = 13, R^2^ = 0.986; F = 7.71 pN, N = 11, R^2^ = 0.986; F = 9.9 pN, N = 22, R^2^ = 0.989). (d) Corresponding half-life values with single exponential fit to the Bell model (blue line). The fit results in t_0_ of 149 s and x_b_ of 0.58 nm (R^2^ = 0.999).

We applied this technique to the binding of complement receptor 2 (CR2) to anti-CR2 antibodies. The anti-CR2 antibodies were coated onto Dynabeads and the CR2 was covalently attached to 3.4 kbp DNA tethers, which also incorporated biotin molecules at the opposite end. The CR2 tethers and the anti-CR2 microbeads were pre-incubated and bound to the imaging chamber surface through biotin-streptavidin interactions (Fig. 4a). After five seconds of recording, the desired force was applied by moving the magnetic tweezers into position. Microbeads would extend their DNA tethers in response to the applied force, followed (in most cases) by a dissociation event, marked by the microbead disappearing from the field of view (Fig. 4b). Datasets lasting for 30 minutes were recorded for three different applied forces: 3.7 pN, 7.1 pN and 9.9 pN. A typical dataset would have several tens of microbeads for analysis.

HoloMiP was used to reconstruct the 3D positions of each microbead through time. Subsequent analysis determined the height above the sample chamber floor that each microbead reached prior to dissociation, and the time of dissociation (Fig. 4b). Our analysis showed that only a subset of microbeads reached full tether extension (1.15 μm) in an immediate response to the applied force (Supplementary Fig. 5c–e). A number of other microbeads moved to some height lower than full tether extension before dissociating. Some other microbeads moved in z through time in a clear stepwise manner, indicating the presence of multiple tether-bead interactions that were dissociating one-by-one (Supplementary Fig. 5b).

To investigate the dependence of the CR2–anti-CR2 dissociation on mechanical load, we considered only microbeads that responded immediately to the applied force and that extended cleanly to at least 1.0 μm (87% of the theoretical full tether extension). This value was chosen as a compromise between the likelihood of a single-molecule interaction and sufficient numbers of microbeads for analysis. Bead dissociation through time was plotted for each applied force and bond lifetimes derived from the fit of single-phase exponential decays to the data (Fig. 4c). The bond half-life values (t_F_) as a function of applied force (F) can then be calculated, yielding data that fit Bell’s equation for slip bonds (28):

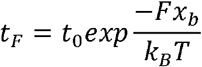

where t_0_ is the zero-force bond half-life, x_b_ is the reaction coordinate, k_B_ is Boltzmann’s constant and T is absolute temperature (Fig. 4d). The results of this analysis indicated a t_0_ value of 149 s and x_b_ of 0.58 nm. These values are in the range of the expected half-lives of antibody-antigen bonds and agree with the mechanical strength of similar antibody-antigen interactions measured by atomic force microscopy (29, 30) (Fig. 4d).

By contrast, the same analysis could not be performed as well on the subset of beads that only extended to a height less than 1.0 μm. Single-phase exponential decays could only be fitted by including a substantial vertical offset term, to account for the number of microbeads that remained tethered at the end of data acquisition (after 30 minutes), indicating the presence of multiple tethers or other long-lived non-specific interactions (Supplementary Fig. 5d). Thus, fast measurement of bead height before dissociation improves the quality of the force spectroscopy data.

## DISCUSSION

We have developed a novel 3D imaging technique we term HoloMiP and applied it to perform force-mediated single-molecule dissociation experiments using magnetic tweezers.

HoloMiP uses a single recorded image to reconstruct the 3D positions of microscopic objects in the sample. Compared to conventional MT techniques, HoloMiP does not require any *a priori* stage movement nor engagement of the magnetic field. Thus, it is ideally suited to measure fast dissociation events. HoloMiP is also superior to conventional look-up table cross-correlation techniques in localising greater densities of objects in 3D.

Whilst the demands on microscope hardware are less onerous for HoloMiP compared to other MT setups, its requirements for computing resources are higher. The holographic reconstruction is computationally intensive; however, this could be mitigated through use of cluster servers or graphics processing units. The LED illumination used limits the effective z range of reconstruction through being relatively low in intensity and coherence length. Although not necessary for the application we demonstrate here, laser illumination would overcome these limitations to provide a greater depth of field for 3D tracking applications.

Force is increasingly recognised as an important factor in biological processes (31, 32). Measuring the dissociation of receptor-ligand complexes under mechanical stress can thus reveal hidden characteristics of the bonds that are relevant to biology (33). As an example, we show the slip-bond character of an anti-CR2 antibody binding to CR2 with mechanical strength typical for affinity-matured antibodies (29, 30). However, with existing single-molecule techniques it is difficult and time-consuming to acquire enough data for new insights into force-mediated single-molecule interactions. HoloMiP is a new technique that can increase the throughput of single-molecule MT force spectroscopy studies. Taking advantage of MT’s virtually uniform magnetic field strength across the microscopic field of view containing many magnetic beads (34), HoloMiP’s ability to track many microbeads simultaneously – including those with overlapping diffraction patterns – means single-molecule force assays can be multiplexed for higher throughput than existing techniques. In addition, HoloMiP overcomes previous limitations of non-specific surface interactions obscuring the single-molecule behaviour by using DNA tethers to separate the interaction of interest from the surface, and through instantaneous measurement of the tether extension under applied force. Furthermore, HoloMiP circumvents a pre-calibration step to acquire a look-up table, which already requires application of a force to the system. Even a small force may disrupt weak single-molecule interactions. Thus, with HoloMiP single-molecule interactions with shorter lifetimes can be investigated. Finally, the localization precision along the z-direction for data analyzed with HoloMiP is more consistent over a larger defocus distance compared to the look-up table method. This opens up MT experiments to a much greater focal range than currently employed. We also note that this means HoloMiP could be adapted easily to other 3D imaging applications, for instance, quantifying fluid flows in microfluidic devices, and tracking the behaviour of free-swimming cells, microorganisms, and other objects.

## ACKNOWLEDGEMENTS

We thank Justin Molloy, Daniel Burnham and Hassan Yardimci for providing starting materials for the DNA tethers and for help with the cross-correlation look-up particle localization, and Antonio Casal for the help with protein conjugation. This work was supported by the European Research Council (Consolidator Grant 648228) and the Francis Crick Institute, which receives its core funding from Cancer Research UK (FC001185), the UK Medical Research Council (FC001185), and the Wellcome Trust (FC001185). This research was funded in whole, or in part, by the Wellcome Trust (Grant number FC001185). For the purpose of Open Access, the author has applied a CC BY public copyright license to any Author Accepted Manuscript version arising from this submission.

## AUTHOR CONTRIBUTIONS

JLF designed and implemented HoloMiP, validated it on synthetic diffraction images and MT data and co-wrote the manuscript. SM implemented look-up table bead localisation and carried out MT calibration and CR2-anti-CR2 dissociation experiments. ILG designed parabolic masking for 3D bead localisation and co-wrote the manuscript. PT conceived and supervised the research and co-wrote the manuscript.

## METHODS

### MT apparatus

We used an imaging system (Cairn Research Ltd) based on an inverted scientific microscope (Ti-E Eclipse; Nikon Instruments Inc.) and fitted with oil immersion objectives (CFI60 Plan Fluor 40×, N.A. 1.30; and CFI Plan Apo TIRF, 100× Oil N.A. 1.49) and automated three-axis stage controller (Applied Scientific Instrumentation Inc.). Semi-coherent illumination was provided by a light-emitting diode (λ = 625 nm; Thorlabs Ltd) with a single lens. This configuration allowed a sufficiently large working distance between the illumination source and the sample for the magnetic tweezer rig to operate. A pair of 5 mm cube neodymium-iron-boron magnets (Supermagnete W-05-N50-G; Webcraft GmbH) was attached to a custom-built bracket that suspended the magnets above the sample, but below the light-emitting diode. The illumination passed between the magnets to the sample. These magnets produced a magnetic field gradient in the sample, and the distance between the magnets was able to be adjusted to modify the strength of the magnetic field gradient. The vertical position of the magnets was adjusted by a voice coil actuator and controller (V-277.630 and C-413.2GA Physik Instrumente GmbH). The voice coil actuator could displace the magnets over the full range of motion of 15 mm in about 100 ms. Digital images were recorded by a sCMOS camera (Orca Flash 4.0 v2; Hamamatsu Photonics K.K.) and passed to a computer for post-processing. Image acquisition was facilitated by MetaMorph software (Molecular Devices LLC).

### Flow cells

Flow cells for use in the MT apparatus were made by sandwiching double-sided tape between two microscope coverslips. The floor surface of the flow cell was cleaned by sonication in solutions of ethanol (95%) and 1 M potassium hydroxide. Silica microbeads (3 μm diameter; Bangs Laboratories Inc.) were partially melted onto the surface to create reference beads, which could be tracked to monitor and correct stage drift. The glass was then treated with silane (2% v/v solution of 3-aminopropyltriethoxysilane in acetone), which was then cured at 110°C for 1 hour. This coverslip was then functionalised with a layer of biotinylated polyethylene glycol (MPEG-SVA and Biotin-PEG-SVA; MW = 5,000; Laysan Bio Inc.). This functionalisation allowed us to attach DNA tethers to the floor of the chamber via biotin-streptavidin bonds

### DNA tethers

A tether of length 3.4 kbp (1.15 μm) was made from a single piece of DNA prepared by polymerase chain reaction (PCR) from a 9.8 kbp pCerOriD plasmid (gift from J. Molloy, Francis Crick Institute).

One primer contained four biotin molecules, which anchored the tether to the floor of the flow cell via biotin-streptavidin bonds. The other primer had functional groups for attachment to the antigen of interest (-thiol, -Digoxygenin). Both were from Integrated DNA Technologies Ltd.

### Conjugation of CR2 protein to DNA tethers

For dissociation measurements, CR2 protein (Bio-Techne) was conjugated to the thiol-modified DNA tethers using sulfo-SMCC (Pierce). Briefly, the tethers were buffer-exchanged into conjugating buffer (PBS pH 7.4, 1 mM EDTA) using desalting columns (Pierce) and incubated with 1M DTT for 30 min at room temperature. CR2 was exchanged into conjugation buffer and incubated with 2.5 mM of Sulfo-SMCC (Pierce), with agitation for 30 minutes at room temperature. The tether and CR2 were then buffer-exchanged into fresh conjugation buffer, mixed and incubated for 1 hour at room temperature. Finally, the sample was buffer-exchanged into PBS.

### Antibodies and conjugation to magnetic beads

Antibodies specific for CR2 (clone b-ly4, BD Biosciences) or for digoxigenin (sheep anti-digoxigenin, Bio-Rad Laboratories) were conjugated to 2.8 μm diameter superparamagnetic Dynabeads (M270 Epoxy; Thermo Fisher Scientific Inc.) using the manufacturer’s protocol. Briefly, 5 mg of beads were resuspended in 1 ml of 0.1 M sodium phosphate buffer pH 7.4, incubated with tilt rotation for 10 minutes and washed once in this buffer. 100 μg of the antibody was buffer-exchanged into 0.1 M sodium phosphate buffer pH 7.4. Equal volumes of the beads, the antibody and 3 M ammonium sulphate in 0.1 M sodium phosphate buffer pH 7.4 were incubated together for 16-24 hours at 37⍰C with tilt rotation. The beads were then washed twice in PBS containing 0.1% (w/v) BSA.

### Force calibration

In our MT apparatus, the force applied to superparamagnetic beads in the sample chamber is a function of magnet height above the sample. This relationship can be deduced by observing the Brownian motion of DNA-tethered beads under the influence of the magnetic field gradient and applying the expression: 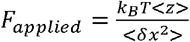, where k_B_ is the Boltzmann constant, T the absolute temperature, <z> the extension of the DNA tether along the optical axis, and <δx^2^> the variance of the bead position (27).

To calibrate the MT system, we used a 48 kbp length of λ-DNA with biotin molecules incorporated into one end, and digoxigenin into the other. The DNA tether attached at one end to the flow cell floor via biotin-streptavidin bonds, and at the other to a Dynabead coated in anti-digoxigenin antibodies. We found our system can apply forces of up to 15.9 pN.

### HoloMiP

The bespoke image processing and object localisation routines of HoloMiP were written in Matlab (R2018a; The MathWorks Inc.). The HoloMiP routines can be sent to multiple CPU cores for parallel processing for shorter reconstruction times. The software is available at https://github.com/iflew/HoloMiP.

HoloMiP first requires relevant physical parameters for the holographic reconstruction as input; namely: wavelength of illumination, refractive index of medium, effective pixel spacing, and z-range and resolution for reconstruction.

Additional processing parameters optimise 3D localisation of objects in the holographic reconstruction: the intensity threshold over which an initial candidate is identified in the image plane; an option for a Gaussian low-pass filter of the image to reduce high-frequency noise; and the dimensions of the cuboids deployed around each initial guess for the x,y,z position of an object candidate, inside which the parabolic masking subroutine runs.

HoloMiP then performs a back-propagation of the illumination field using the Rayleigh-Sommerfeld formalism (17, 23) at each specified z position to result in a 3D volume of electromagnetic field intensities (Fig 1b,c).

A hologram (*I_H_*(*x,y*, 0)) results from the interference of an object term, *Õ*(*x,y*, 0), which arises from light scattered off the sample, with an unscattered reference term, 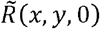:

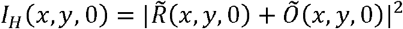

where z=0 is denoted the hologram plane (focal plane).

For our purposes, a separate recording of a background image can be used as a good approximation of the intensity of the reference term: 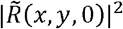

To reconstruct the electromagnetic field *Õ*′(*x, y, z*) at a given position, z, along the optical axis, we convolve the recorded hologram object term *Õ*(*x,y*, 0) with the Rayleigh-Sommerfeld propagator:

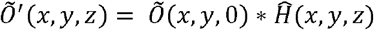

where 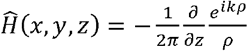.
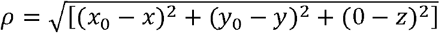, the distance between a given coordinate in the hologram (*x*_0_*y*_0_,0) and a reconstructed position (*x,y,z*). 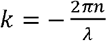, the wavenumber of light with wavelength *λ* and in a sample medium of refractive index *n*.

HoloMiP performs this operation in Fourier space, using 2D Fast Fourier Transforms:

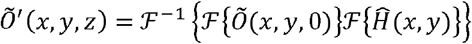

The use of the Fourier transform allows for a low-pass Gaussian filter to be applied to the recorded hologram, if desired, to smooth high-spatial frequency noise, with minimal additional computational overhead.

The procedure is repeated for each user-defined z value, resulting in a 3D electromagnetic field representing the recorded sample. The square of this field is then taken to provide an 3D intensity map (Fig. 1c).

### Object localisation

The next phase of the HoloMiP algorithm is to localise in 3D space the objects in the sample.

Adapting the approach taken in (24) we find identifying features in the gradient of the intensity field more reliable than working on the intensity field itself. We compute the gradient along the z direction by applying a Sobel-like kernel:

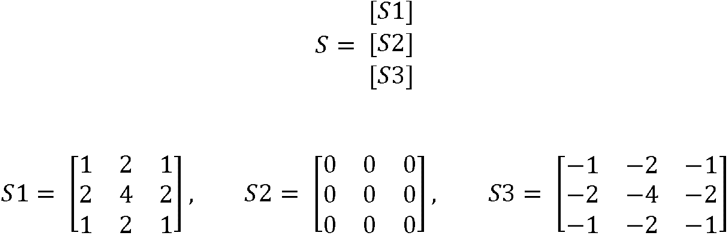

Supplementary figure 1 demonstrates the effect of applying this gradient kernel to the 3D intensity field and supplementary figure 2 demonstrates the effect the kernel application has on microbead localisation along the optical axis.

Application of the kernel is computationally expensive. Thus, to minimise the volume over which the kernel is applied, we first localise the objects of interest in the hologram plane and extract columnar cuboids (typically 15 pixels across) centred on these values (Fig. 1e). The initial x, y localisation uses a 2D peak-finding algorithm on a maximum-pixel projection of the reconstruction volume (Fig. 1d). At this stage object candidates can be excluded based on their proximity to other objects or the edge of the hologram, if desired.

After application of the Sobel-like kernel (Fig. 1f), an approximate z-value for an object position within the columnar sub-volume is determined by a 1D peak-finding algorithm applied to the central line of pixels along the z-axis (Supplementary Fig. 2).

A cuboid sub-volume, typically 15 × 15 × 25 pixels (Fig. 1g) centred on the approximate x, y, z values determined as described above is then passed to a subroutine for parabolic masking (Fig. 1h,i). The voxel intensity values within this cuboid are modelled by a paraboloid, which we use as a proxy for object position (25). Thus, to localise precisely the centre of this paraboloid, we:

1. Multiply the sub-volume by a 3D paraboloid of identical size and determine the centre of this product by summing voxel intensities along each dimension;
2. Determine the difference between the initial and new centre positions;
3. Repeat this process until this difference converges to a user-defined threshold;

The precise 3D positions of all objects in all frames are then stored for subsequent analysis (Fig 1.j). For instance, the 3D position of each object can be tracked through time and correlated to the introduction of the magnetic field gradient from the MT. In this way dissociation events can be observed, and dissociation times measured. The extension length of each tethered microbead can be computed also to ensure complete tether extension, from which single-molecule interactions can be inferred.

### Computer hardware and processing speeds

For object 3D localisation with HoloMiP we used an Asus workstation (AsusTek Computer Inc.) with an Intel i9 CPU with 14 cores (i9-7940X CPU; 3.10 GHz; Intel Corporation) and with 128 GB total RAM running Microsoft Windows 10 Enterprise 2016.

Holographic reconstruction is potentially computationally expensive given the iterative nature of applying the propagator to the hologram to build up the 3D volume. In addition, the requirement to hold the reconstructed 3D volume in memory to perform object localisation sets conditions on computer RAM.

To minimise computation time, we employed the multiple CPU cores, such that up to 14 frames (of size 1048 px × 1048 px × 100 z-slices) can be reconstructed in parallel using Matlab’s parfor command. To minimise the burden on computer RAM, we designed HoloMiP to perform localisation operations on small sub-volumes around each identified object candidate (Fig. 1e-i).

Compared to single core computing, our strategy of parallel reconstruction is slower for some of HoloMiP’s operations, owing to the additional time taken to transfer data onto different cores. However, this is only apparent for processing of very small numbers (<20) of frames. Over 200 frames, operating HoloMiP on parallel cores results in at least a four-fold reduction in processing time compared to single CPU operation.

It is important to note that the 3D localisation of objects via parabolic masking is the least computationally expensive part of the routine (taking 500-1000 times less time than 3D reconstruction of the volume from the hologram, for instance). Thus, for parallel core operation, the total computational time of deploying HoloMiP is almost independent of the number of objects in the field of view. This is not the case in most other comparable 3D object localisation implementations, where each object must be localised sequentially in each frame in a dataset.

On our workstation, a hologram of size 1048 px × 1048 px and reconstructed to 100 z-slices will return 3D positions of all objects in around 1.1 seconds.

We note that significant reductions in processing times could be made by:

- implementing HoloMiP in a performance-oriented language;
- using graphics processing units, which are better suited to iterative computation, to perform holographic reconstruction; or
- implementing HoloMiP on high-performance computer clusters.

### Synthetic data

We generated synthetic diffraction patterns simulating microbeads *in silico* to compare the performance of HoloMiP and the look-up table cross-correlation technique as follows.

A tethered microbead was held above the sample surface by applying the magnetic tweezers. The microscope stage was moved along the optical axis and a z-stack of images with a 50 nm spacing was captured. This image stack was converted into a look-up table consisting of a stack of radial diffraction profiles as a function of bead distance from focus, using the techniques described in (16).

To generate a synthetic bead diffraction image, a single profile was selected from this look-up table, revolved on a polar grid and then placed on a larger pixel grid (1024 px × 1024 px) at the specified location. Multiple diffraction patterns can be generated from different radial profiles and placed where desired. Setting the intensity of the diffraction patterns to be centred on zero ensured overlapping patterns summed appropriately. Poisson-distributed noise was added to simulate camera shot noise. The intensity of the image was rescaled to the range of the pixel values recorded by our microscope camera. The resulting image, or set of images, was then sent to HoloMiP or lookup table cross-correlation algorithms for 3D localisation.

