## Supplementary Figures for "Digital holography-based 3D particle localisation for single molecule tweezer techniques"

Supplementary Material

**James L. Flewellen^1,2^, Sophie Minoughan^1^, Isabel Llorente Garcia^3^, Pavel Tolar^1,2,^ ***

^1)^ Immune Receptor Activation Laboratory, The Francis Crick Institute, London NW1 1AT, United Kingdom

^2)^ Institute of Immunity and Transplantation, Division of Infection and Immunity, University College London, London, NW3 2PP, United Kingdom

^3)^ Dept. of Physics and Astronomy, University College London, London, WC1E 6BT, United Kingdom

**Optical correction for retrieval of absolute particle positions**

To determine the optical correction required for 3D imaging with microscope objectives, and to determine the optimal focal range for experiments, a sample was prepared with Dynabeads and silica reference beads immobilised on the imaging chamber floor. The focus of a 40x objective lens was initially positioned at the floor of the chamber. Using a piezoelectric nano-positioner, the objective lens was moved in 0.5 µm steps away from the floor of the chamber over a range of 12 µm. The z positions of the microbeads were then recovered using HoloMiP and compared to the theoretical position of the objective (Supplementary Fig. 3a). We observed a linear relationship between recovered z position and objective position when the objective was positioned between 7 µm and 12 µm from the chamber floor (Supplementary Fig. 3b).

To validate this result further, and to simulate a typical tether extension experiment, we used the same sample and moved the objective away from its focus at the floor of the sample by 12 µm. The imaging stage was then moved over 5 µm in 100 nm steps, bringing the sample closer to the focus (Supplementary Fig. 3c). The z positions were again determined by HoloMiP and compared to the theoretical sample-to-focus distance. The linear relationship between recovered and theoretical position held for both Dynabeads (Supplementary Fig. 3d) and silica reference microbeads (Supplementary Fig. 3e).

It is important to note that while the absolute position in 3D space of the centre of the microbead is not recovered, its position in 3D *relative* to other beads in the sample, and to itself at different time points in a recording is, through the reconstruction of light scattered off the bead using HoloMiP. The electric field distribution of microspheres under similar imaging conditions is discussed in (1). Operating within the linear focus regime of the objective lens ensures that comparison of relative positions of microbeads is trivial. It is also possible to apply a non-linear correction to samples recorded outside of this range, as long as this relationship has been determined.

**Microbead position offset between HoloMiP and LUT approaches**

There is a small lateral offset when comparing the recovered positions found using HoloMiP and the conventional look-up table approach. The magnitude of this offset increases linearly with the microbead distance away from the objective focus (Supplementary Fig. 4). We thus posit that HoloMiP is more sensitive to the alignment of the optical axis to the camera sensor. To quantify this offset, we used the two techniques to analyse a force calibration dataset, where a Dynabead is tethered to a 16.3 µm length of λ-DNA and subject to an increasing magnetic field. This dataset yields 40,000 images of a Dynabead at a range of displacements from the microscope focus. The difference in the output of the two techniques was computed and a linear fit applied along both x and y axes to determine the lateral offset as a function of microbead z position. The difference between the two techniques was found to vary by 17.19 ± 0.03 nm along x and 22.09 ± 0.03 nm along y per 1 µm displacement from the focus (uncertainty is 95% confidence bound on the linear fit). This correction factor can be used to make the two techniques to agree with each other.

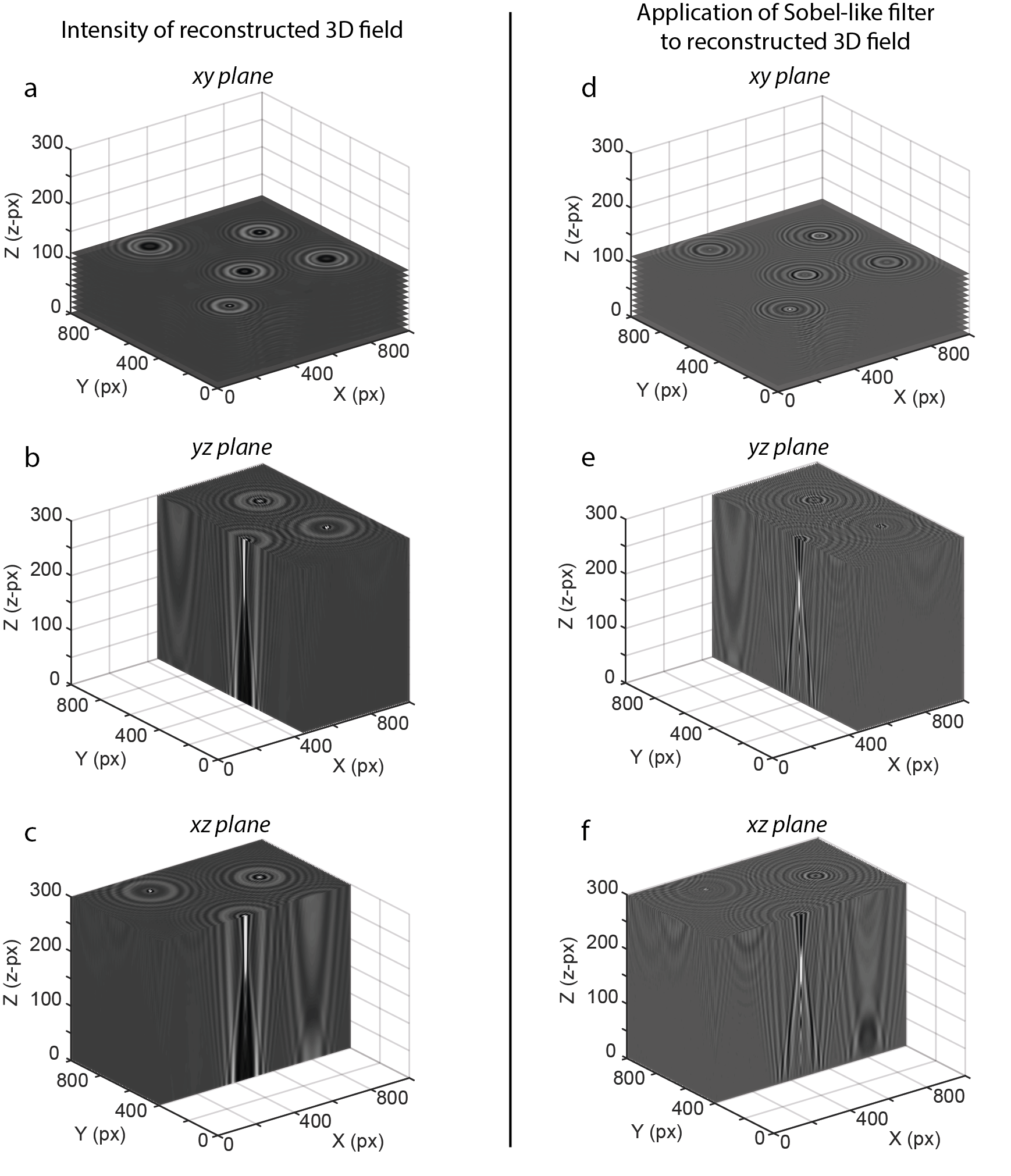

**Supplementary Figure 1.** Holographic reconstruction and the effect of applying a 3D Sobel-like filter. The intensity of the back-propagated 3D electromagnetic field from the hologram shown in Fig. 1c is displayed in (a), (b) and (c) with cutaways along different planes. (d), (e) and (f) show the same cutaways after the Sobel-like filter has been applied to the data, resulting in the gradient of the field along z. Features in the 3D field, such as the bright spots corresponding to the positions of microbeads, are now easier to detect. Supplementary videos 1 to 6 show animations scrolling through these reconstructed volumes. Pixels along x and y are 0.065 µm; each pixel value along z corresponds to 0.1 µm.

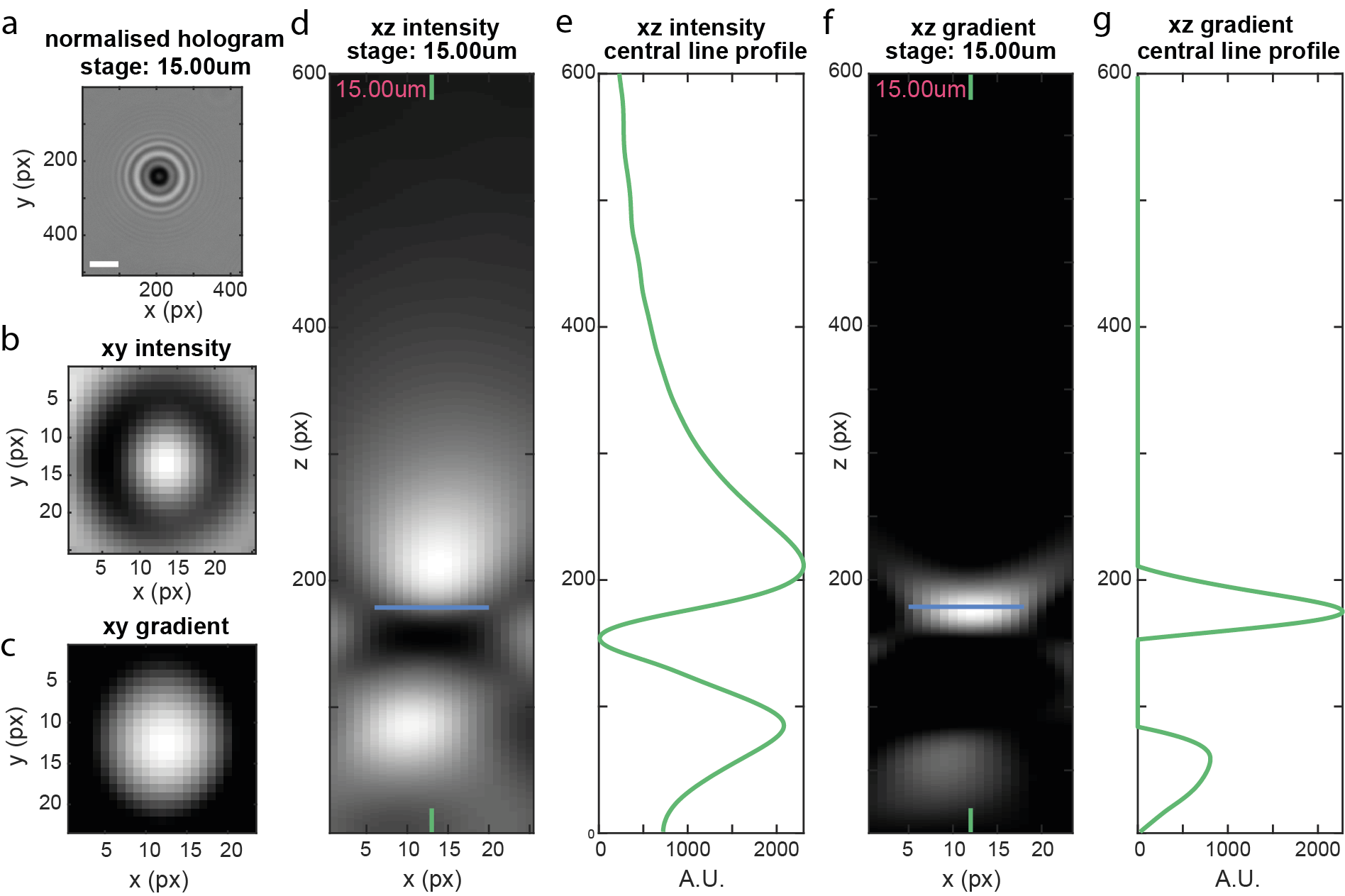

**Supplementary Figure 2.** Application of 3D Sobel-like filter to holographic reconstruction. (a) Recorded normalised hologram, of a 2.8 µm-diameter Dynabead. The stage is positioned 15 µm below the focal plane. The scale bar is 5 µm. (b) Zoom-in of the intensity of the holographic reconstruction *at the z-value of initial object detection* (blue horizontal lines in (d) and (f)). (c) Same plane as in (b) but after the Sobel-like gradient filter has been applied. (d),(f) xz-planes through the centre of the microsphere from the focal plane (1) to the extent of the holographic reconstruction (600). (d) Intensity of the reconstructed field; (f) the field after the Sobel-like gradient filter has been applied. (e), (g) Vertical line profiles through the centres of (d) and (f), respectively. The taller peak of the trace in (g) is used as a proxy for position of the microbead along z. The value of this peak (blue line in (d) and (f)) is the initial guess fed into the 3D parabolic masking algorithm for more precise localisation. Supplementary video 7 shows an animation of these data scrolling through all reconstructed planes to a distance of 35 µm from the focus. Pixels along x and y are 0.065 µm wide; each pixel value along z corresponds to 0.1 µm. A.U. = arbitrary units.

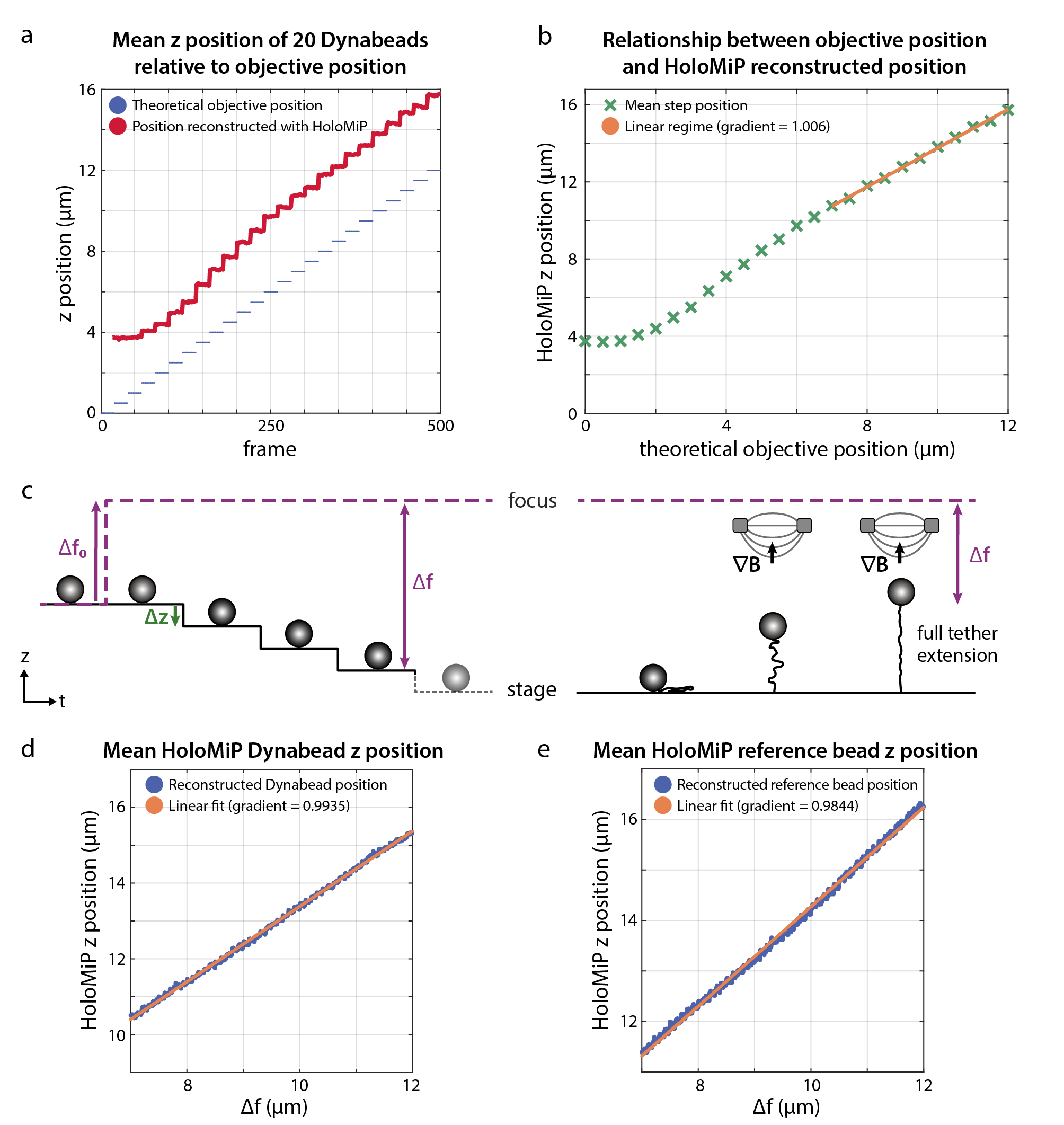

**Supplementary Figure 3.** Optical correction for 3D imaging with 40x objective. (a) After focusing on the floor of a sample of immobilised microbeads, the objective lens is moved in discrete steps (blue). The 3D positions of beads are recovered using HoloMiP (red). (b) We observe a linear relationship between objective position and z position determined by HoloMiP when the objective is positioned between 7 and 12 µm from the sample. (c) An experiment to validate this result begins by moving the objective focus a distance Δf_0_ from the sample chamber floor. The stage is then moved away from the focus in discrete steps Δz. This simulates the behaviour of a Dynabead in a magnetic tweezers experiment (right). We find the linear relationship between sample-to-focus distance (Δf) and z position recovered using HoloMiP holds for the range of 7 to 12 µm for both Dynabeads (d) and silica reference beads (e).

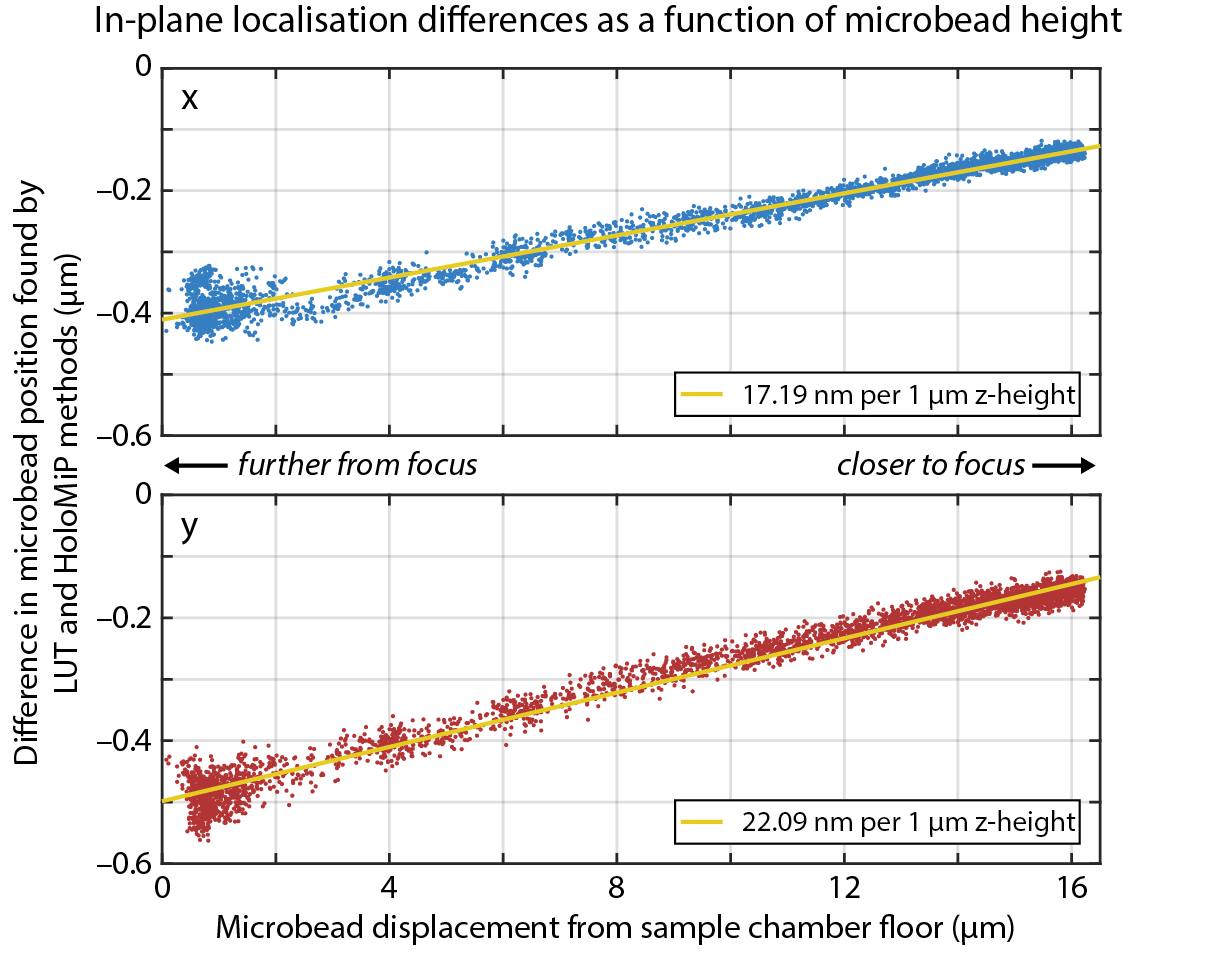

**Supplementary Figure 4.** The 3D position of a 2.8 µm-diameter Dynabead tethered to a 16.3 µm DNA tether in response to an increasing magnetic field was determined by both the look-up table cross-correlation method (LUT) and HoloMiP. The difference between these two techniques shows an offset along both the x (top) and y (bottom) axes, which correlates to a linear function (yellow line). As stated in the text, we conclude this offset is a function of the way in which the LUT technique averages a 2D diffraction pattern to produce a 1D radial profile, thus imposing an artificial radial symmetry on the ring patterns and rendering the technique insensitive to minute differences in angle between incident light and the normal to the imaging plane. The holographic reconstruction in HoloMiP does not involve radial averaging. We determined an angle between optical axis and normal to the imaging plane of around 1.1°.

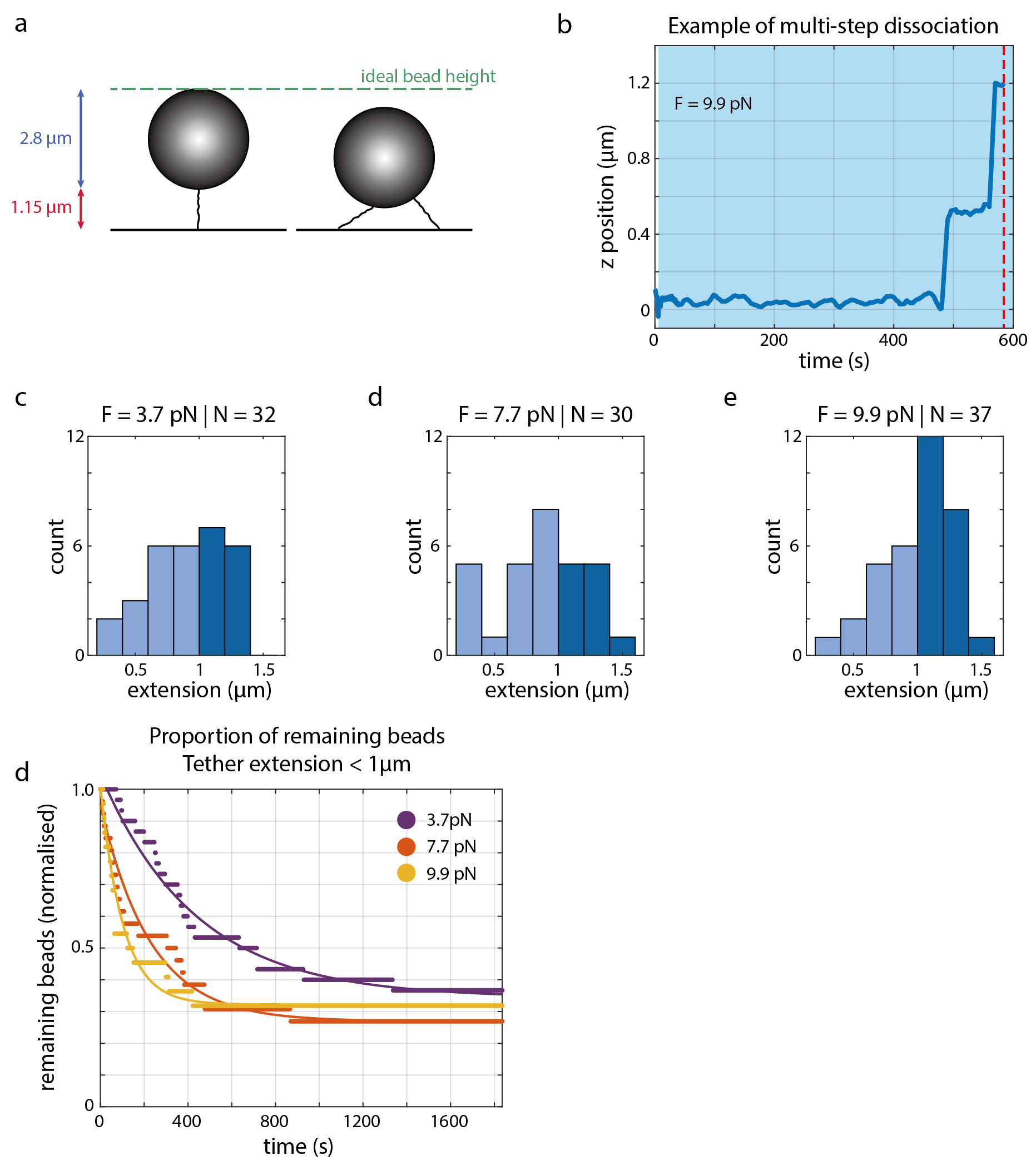

**Supplementary Figure 5.** Evidence of multiple tether attachments in MT experiments. (a) Scale diagram showing Dynabeads with multiple tether attachments will not extend to the ideal bead height in magnetic tweezer experiments. (b) Example z position trace of a Dynabead exhibiting multi-step dissociation in CR2–anti-CR2 force-mediated unbinding experiments. The blue shading indicates the applied force; the red dashed line indicates when the microbead fully dissociates. (c, d, e) Histograms for each applied force showing the extensions that tethered microbeads reached before dissociation. We selected microbeads that reached 1 µm extension (dark shading) for the analysis shown in Fig. 4. The remaining beads were analysed in (d) and (e). Beads exhibiting multi-step behaviour were excluded. (d) Proportion of microbeads with a tether extension < 1 µm in the CR2–anti-CR2 dataset remaining attached through time. Single exponentials could only be fitted with a high plateau term to account for the large proportion of microbeads that did not dissociate by the end of the recording (F = 3.7 pN, N = 19 with 11 remaining at end of recording, R^2^ = 0.989; F = 7.7 pN, N = 19 with 7 remaining at end, R^2^ = 0.985; F = 9.9 pN, N = 15 with 7 remaining at end, R^2^ = 0.985).

**Supplementary video 1.** xy planes of 3D intensity field reconstruction (related to Supplementary Fig. 1a).

**Supplementary video 2.** yz planes of 3D intensity field reconstruction (related to Supplementary Fig. 1b).

**Supplementary video 3.** xz planes of 3D intensity field reconstruction (related to Supplementary Fig. 1c).

**Supplementary video 4.** xy planes of 3D gradient field reconstruction (related to Supplementary Fig. 1d).

**Supplementary video 5.** yz planes of 3D gradient field reconstruction (related to Supplementary Fig. 1e).

**Supplementary video 6.** xz planes of 3D gradient field reconstruction (related to Supplementary Fig. 1f).

**Supplementary video 7.** Animation showing effect of Sobel-like gradient filter (related to Supplementary Fig. 2).
